# Selective approach behavior toward context-dependent ultrasonic vocalizations in male mice

**DOI:** 10.64898/2026.04.10.717606

**Authors:** Katsumasa Takahashi, Kazuma Hase, Takumi Miyajima, Jumpei Matsumoto, Tetsufumi Ito

**Author notes:** **Corresponding authors**: Katsumasa Takahashi,; Tetsufumi Ito.

## Abstract

Ultrasonic vocalizations (USVs) are widely used in rodent social communication, yet the functional significance of male–male vocal interactions in mice remains unclear. Here, we investigated whether USVs produced during specific social behaviors influence the behavior of conspecifics. Using playback experiments, we compared responses to vocalizations recorded during chasing and being chased in male–male interactions. We found that USVs emitted by chased intruders consistently elicited approach behavior in receiver mice, whereas those emitted by chasing individuals did not. Acoustic analyses revealed that these vocalizations differed in syllable composition, with intruder calls containing a higher proportion of upward frequency-modulated syllables and exhibiting higher mean frequencies. In addition, the temporal organization of syllables appeared to contribute to the behavioral response. Together, these results suggest that male mice respond selectively to certain USV patterns associated with specific social contexts, indicating that acoustic features and temporal structure may jointly influence social approach behavior in mice.

**Highlights:**

- Behavioral context (chased vs. chasing) shapes the composition of USV syllable types
- Male mice selectively approach USVs from chased intruders, but not chasing residents
- The approach response exhibits high temporal synchrony across individual receivers
- Temporal organization of syllables modulates approach behavior based on acoustic features

## 1. Introduction

Ultrasonic vocalizations (USVs) are one of the key modalities for rodents to communicate with conspecifics. The most well-known example is the “pup calls” by neonatal rodents when isolated from their mothers, a phenomenon observed across species, including mice ^1^, rats ^2^, and gerbils ^3^. These vocalizations, which are triggered by anxiety or decreased body temperature, play a critical role in mother–infant interactions, as they can elicit maternal retrieval of the pup to the nest. In rats, it is well established that 50 kHz USVs are associated with positive affective states, such as those induced by tickling ^4^ or administration of cocaine ^5^, whereas 22 kHz USVs are typically linked to negative affective states elicited by aversive stimuli, including predator odor ^6^ or pain ^7^. Playback experiments using 50 kHz and 22 kHz USVs have been shown to elicit approach and defensive behaviors, respectively ^8,9^. These findings suggest that USVs influence the behavior of other individuals.

Importantly, in mice, not only the “pup calls” but also the “male song” produced by adult males in the presence of females have been reported to modulate the behavior of conspecifics. The “male song” is considered to serve as a courtship signal and consists of phrases composed of several types of syllables that are temporally organized and repeated ^10^. Previous studies have shown that both types of USVs induce approach behavior in female mice, while not affecting males ^11–14^, with the exception that father males exhibit approach behavior specifically toward pup calls ^15^. This suggests that male mice typically do not perceive these specific vocalizations as signals relevant to themselves.

However, male mice also emit USVs during male-male interactions, such as in the resident-intruder paradigm ^16^. The intrasexual vocal interactions in rodents have been studied less intensely than the vocal interactions cited above because of the difficulty in identifying the individual source of these vocalizations. Recent advancements in multi-microphone sound-source localization have begun to overcome this limitation ^17–19^. For instance, USV characteristics differ markedly depending on whether a male is chasing or being chased ^20^, implying that vocalizations vary with behavioral context. Nevertheless, the functional impact of these male-to-male USVs on the receiver’s behavior remains poorly understood.

In this study, we investigated how USVs emitted during specific social contexts influence male mouse behavior using playback experiments. We focused on USVs recorded during “chasing” and “being chased” behaviors, which possess distinct acoustic profiles ^20,21^. Our results demonstrate that behavioral responses vary according to the acoustic features of USVs, providing strong evidence that male mice recognize specific USVs as meaningful social signals. These findings enhance our understanding of the complexity of social interactions and vocal communication in rodents.

## 2. Methods

### 2.1. Animals

A total of 46 male CBA/J mice of age ranging from 2 to 6 months were used in this study. The CBA strain was selected for its well-preserved hearing sensitivity into adulthood and does not show impairment of hearing sensitivity before 15 months old ^22^. This is important for this study because this strain lacks early-onset age-related hearing loss in the high-frequency range used in USVs, which are commonly found in other popular strains such as C57BL/6. Eighteen mice were used to record ultrasonic vocalizations during social interactions, while the remaining 28 participated in the playback experiment; however, one mouse was excluded from all subsequent analyses because it escaped from the apparatus during the test session.

All mice were housed in temperature-controlled rooms maintained at 20–24 ℃, with 2–4 animals per cage, under a 12:12 h light/dark cycle (lights on at 7:00 a.m.). All mice were at least 8 weeks old at the start of the experiment and had ad libitum access to food and water. All experimental procedures were conducted in accordance with the Regulations for the Handling of Animal Experiments of the University of Toyama and were approved by the University of Toyama Animal Experiment Committee (A2023-MED01) and the Guidelines for Proper Conduct of Animal Experiments by the Science Council of Japan.

### 2.2. Apparatus

USVs were recorded using the USVCAM system (**Fig. 1A**) ^19^. Briefly, mice were allowed to interact in a rearing cage with two layers of walls, one made of sound-absorbing material (e.g., paper towels) and the other of stainless mesh, inside. A custom cage lid with a clear mesh screen was employed to prevent escape while maintaining visibility. Mouse behavior was monitored using an infrared camera (RealSense L515, Intel, CA, USA), and USVs were recorded with four ultrasound microphones (TYPE 4158N, ACO, Tokyo, Japan) in the sound-attenuating chamber. The audio data was captured by each microphone, amplified using a four-channel microphone amplifier (BSA-CCPMA4-UT20, Katou Acoustics Consultant Office, Kanagawa, Japan), and sampled at 400 kHz with a DAQ device (DT9816-S, Digilent, TX, USA). To identify individuals that emitted USVs, audio and video data were temporally synchronized and stored on a PC, where they were then processed using custom software written in Python (version 3.11.9; Python Software Foundation, NC, USA).

**Figure 1.**
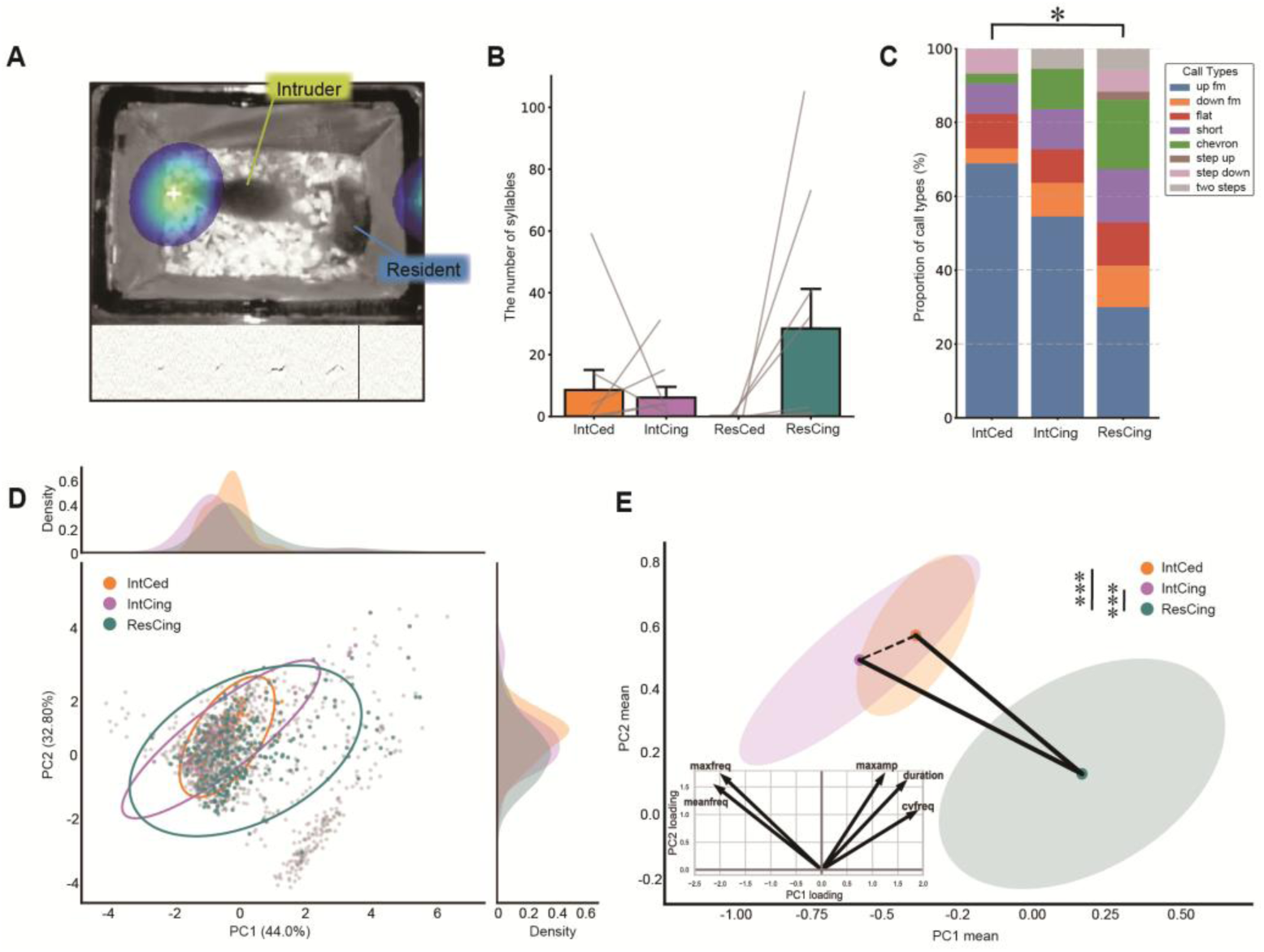
Distinct ultrasonic vocalizations (USVs) produced by chased intruders and chasing residents in male mice. (A) Sound source localization of USVs during a resident-intruder test. (Top) A single video frame processed by USVCAM. The heatmap represents the estimated spatial origin of the emitted USVs. (Bottom) A spectrogram showing the recorded USVs. (B) Number of syllables emitted in 10 minutes. Abbreviations: IntCed, “intruder, being chased”; IntCing, “intruder, chasing”; ResCed, “resident, being chased”; ResCing, “resident, chasing”. (C) Percentage of call types emitted by male mice in each group. *: *p*<0.001, Fisher’s exact test. (D) Scatter plot of syllables emitted in 3 different social contexts [i.e. IntCed (orange), IntCing (purple), and ResCing (green)] on 2 principal component (PC) axes (center), and histograms of syllables on each PC axis after the principal component analysis (PCA) using five acoustic parameters. Ellipses represent the 95% confidence regions estimated from the robust covariance. (E) Bootstrap distribution of the mean PC scores. The colored ellipses indicate the 95% confidence regions for the mean values obtained from bootstrap resampling (n = 10,000 iterations) within each social context. Group pairs connected by solid lines represent significant differences, while those connected by dashed lines indicate no significant difference. Statistical significance was determined using bootstrap-based hypothesis testing of the difference between group means (***: *p* < 0.001).

The behavioral experiment was conducted in a custom-built “I-maze” (**Fig. 2C**; 10 cm wide, 43 cm long, and 26 cm high) installed within a sound-attenuating chamber under infrared illumination. The floor and lateral walls of the maze were constructed from aluminum composite panels, while the end walls were made of stainless-steel mesh to prevent mice from climbing. Speakers (ES1, Tucker-Davis Technologies, FL, USA) connected to sound drivers (ED1, Tucker-Davis Technologies, FL, USA) were positioned outside both ends of the maze, and a video camera (DMK33UX273, The Imaging Source, Bremen, Germany) was mounted above the apparatus. To trigger the presentation of auditory stimuli based on the real-time position of the mouse, data from the camera were processed using custom software written in MATLAB (Mathworks, MA, USA) and output to the sound driver via a DAQ device (USB-6346, National Instruments, TX, USA).

**Figure 2.**
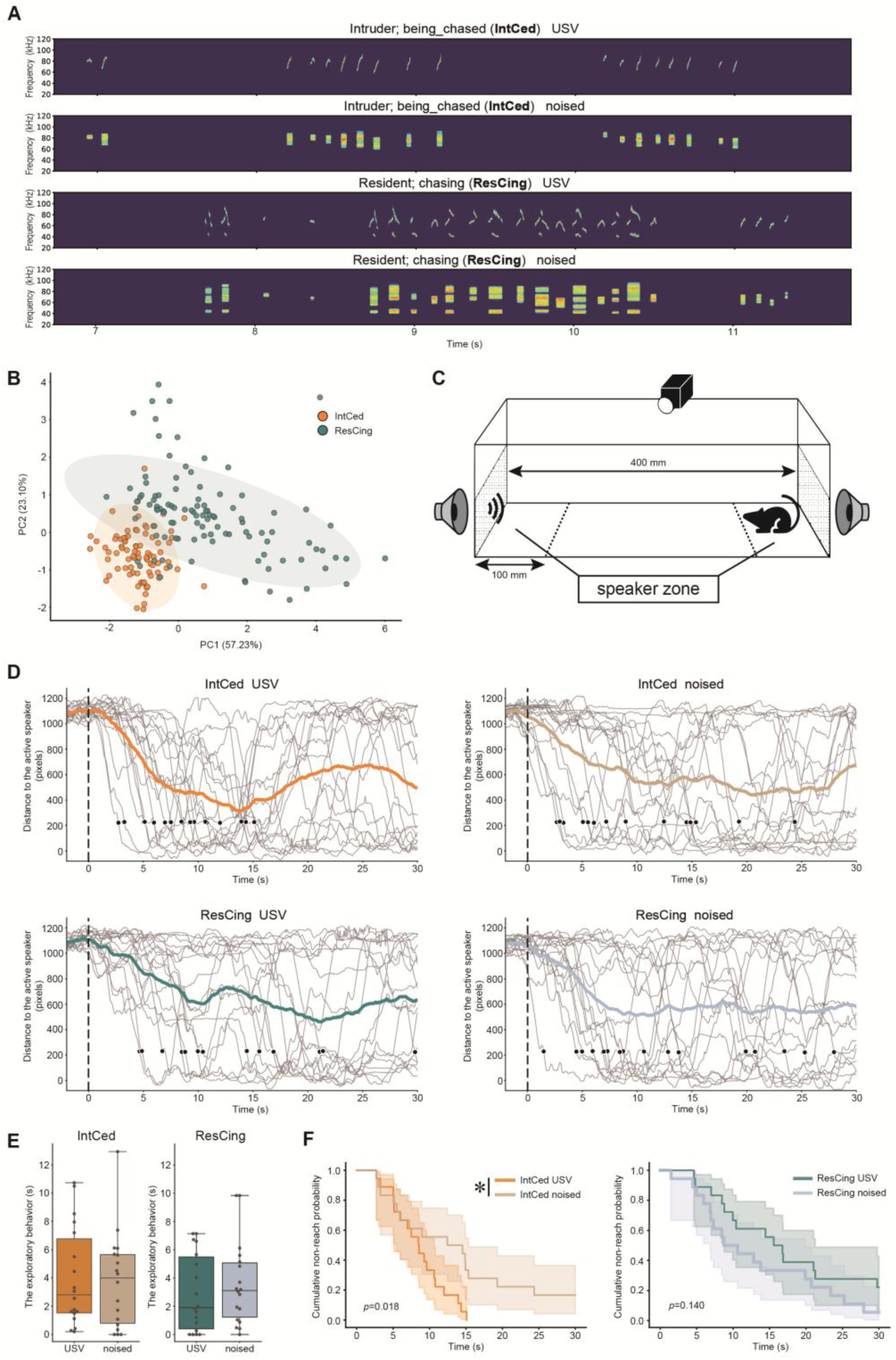
Male mice approach the USVs of chased intruders in a non-habituation context. (A) Representative spectrograms of sound stimuli. USV (top) and their corresponding phase-randomized sound (“noised”, bottom) are shown for both the IntCed and ResCing USVs. (B) Scatter plot of syllables on 2 PCs of the acoustic features of USVs used in the playback experiment. (C) Schematic illustration of the experimental setup. The zones at both ends beyond the dashed line are defined as “speaker zones”. (D) Time course of the distance to the active speaker. Gray lines represent individual data from 18 mice, and bold-colored lines represent the mean. The dashed line indicates the onset of the auditory stimulus, and the black dot marks the first entry of each mouse into the speaker zone. (E) Comparison of exploratory behavior toward the sound source (for definition, see Methods) between USV and noised playback in the IntCed and ResCing groups. (F) Survival analysis of time to enter the active speaker zone. The shaded area indicates 95% CI. *: *p*<0.05, log-rank test.

### 2.3. Procedure

#### 2.3.1. USV Recording and analysis

At least one week before recording, male and female mice were housed together in the same cages that would be used for the recording sessions. Immediately before recording, the female was removed, and the cage containing a single male (referred as “resident” hereafter) was placed inside the sound-attenuating chamber of the USVCAM system for a 5-minute acclimation period. A novel male (referred as “intruder”) was then introduced into the cage, and their social interaction and vocalizations were recorded for 10 minutes.

Mouse interactions were analyzed using video data by assigning timestamps to periods of “chasing” and “being chased,” and audio data corresponding to these periods were subsequently extracted. We focused on the emitter identity (resident [Res] or intruder [Int]) and the direction of movement (chasing [Cing] or being chased [Ced]). USVs were thus classified into four categories: resident chasing the intruder (ResCing), resident being chased by the intruder (ResCed), intruder chasing the resident (IntCing), and intruder being chased by the resident (IntCed). USV syllables obtained from the audio recordings of each behavioral category were analyzed using a slightly modified version of the USVSEG algorithm ^23^, enabling the quantification of syllable count, duration, maximum frequency (maxfreq), maximum amplitude (maxamp), mean frequency (meanfreq), and the coefficient of variation in frequency (cvfreq). In addition, syllables extracted from background noise were collected and saved as a separate dataset for the playback experiment.

#### 2.3.2. Playback experiment

The ResCing and IntCed categories of sound stimuli, which differed significantly more than other categories (**Fig. 1C, E**), were used in the experiment. Each stimulus consisted of a 30-second sequence edited from the audio data of a single mouse, containing USVs representative of each behavioral category, with amplitude normalization applied. For each “USV” stimulus, a corresponding “noised” stimulus was generated as a control, resulting in a total of four sound stimuli presented to the mice (**Fig. 2A**). The “noised” is a sound stimulus in which USV syllables are processed using a fast Fourier transform (FFT), with their phase components randomized while preserving the original amplitude spectrum. This procedure preserves the overall spectral content of the original syllables but disrupts their temporal structure.

##### Experiment 1 – playback in a novel context

Eighteen mice were used. The playback test was conducted during the dark phase, and male mice were allowed to explore the I-maze under dark conditions freely. Infrared illumination enabled real-time detection of the animals’ positions for video recording. Mice were placed in the apparatus for at least 30 seconds before the stimulus began. The sound stimulus (54 ± 3 dB SPL at a distance of 5 cm from the speaker) was presented when a mouse remained spontaneously in the distal zone of the maze (< 10 cm from the distal end, opposite the active speaker) for 2 s. To detect the mice, we applied binarization to identify a region that differed from the background images captured in advance, and used the region’s centroid to estimate the mouse’s position. After the end of each sound stimulus, a subsequent stimulus was presented under the same conditions following a silent interval of at least 30 seconds. The presentation order of the ResCing and IntCed phases, as well as the order of USV and noised conditions within each phase, was counterbalanced across subjects. At the end of each experimental session, the mice were removed from the apparatus, and any remaining feces and urine were cleaned. Then, the apparatus was wiped with 70% ethanol to eliminate odors. Behavior during the test was recorded using an overhead camera, which allowed for the tracking of the mouse’s movements. For body part tracking (head, right ear, left ear, neck, back, and tail base), we used DeepLabCut (version 2.3.9)^24,25^. Specifically, we labeled 140 frames taken from 7 animals. We used a ResNet-50-based neural network with default parameters for 50,000 training iterations. This network was then used to analyze videos from similar experimental settings.

##### Experiment 2 – playback in a familiar context

Nine mice were used. On the day before the playback experiment, mice were allowed to move freely in the dark for 10 min to acclimate them to the I-maze. The playback experiment was then conducted using the same procedure as in “Experiment 1,” but a “silent” experiment without sound stimulation was introduced as an additional experimental condition alongside the USV and noised.

### 2.4. Statistical analysis

All behavioral and acoustic data were analyzed using custom Python and MATLAB scripts. For syllable classification, the recording data was processed in MATLAB using VocalMat ^26^. To characterize acoustic features of USV syllables, principal component analysis (PCA) was performed on the five acoustic parameters extracted using USVSEG (see 2.3.1), for each behavioral category. To compare syllable characteristics across behaviors, a bootstrap test (10,000 iterations) using the PC1 and PC2 score vectors was conducted after downsampling to equalize syllable counts across categories. Fisher’s exact test was performed using R (version 4.2.2; R Foundation for Statistical Computing, Vienna, Austria) to determine whether the distribution of syllable types differed significantly among behavioral contexts. Following the overall test, standardized residuals from the chi-square test were calculated to identify specific categories contributing to the observed associations. Cells with an absolute standardized residual larger than 3 were considered to exhibit a strong deviation from expected frequencies. For the playback experiments, approach behavior was evaluated using Kaplan–Meier curves to illustrate the latency to enter the speaker zone (see **Fig. 2C**). Statistical differences between sound stimuli were assessed using the log-rank test. Exploratory behavior toward the sound source was defined as the mouse being positioned within the speaker zone while facing the sound source (head oriented within ±45° relative to the center of the maze end). To examine the temporal relationship between syllable presentation and approach behavior, Pearson correlation coefficients were calculated using frame-by-frame analyses. For each trial, the distance between the mouse and the sound source and the occurrence of syllables were correlated across all frames during the first half of the stimulus presentation period. To assess consistent timing of approach behavior, the distribution of correlation coefficients for each second was compared against chance level (0) using one-sample t-tests. Statistical significance was set at *p* < 0.05.

## 3. Results

### 3.1. Male mice produce different USVs depending on context and behavior

We recorded USVs from male mice during the resident–intruder task using USVCAM to identify vocalizing individuals (**Fig. 1A**) and to determine which vocalizations were produced in association with specific behaviors. When syllables counted specifically during chasing and being chased behaviors, USVs were observed in all conditions except when resident mice were being chased (ResCed) (**Fig. 1B**). A total of eight distinct call types were identified across the observed behavioral categories (**Supplemental Fig. 1**). Extended Fisher’s exact test with Monte Carlo simulation for 3×8 matrix revealed a significant association between these behavioral categories and call types (*p* < 0.001; **Fig. 1C**). Subsequent *post-hoc* comparisons with Bonferroni correction showed significant differences in syllable distributions, specifically between IntCed and ResCing behaviors (*p* < 0.001; **Fig. 1C**), while no other significant differences were found (IntCed vs IntCing: *p* =0.083, IntCing vs ResCing: *p* = 0.155; **Fig. 1C**). Standardized residual analysis further identified the specific syllables contributing to these differences; “up-fm” and “chevron” syllables exhibited marked deviations from expected frequencies (**Supplemental Table 1**), indicating that the former and latter types are selectively emitted during IntCed and ResCing situations, respectively. Additionally, to visualize and verify the acoustic distinctiveness of the syllables, we performed a PCA based on five acoustic features: duration, maxfreq, maxamp, meanfreq, and cvfreq. The first two principal components (PC1 and PC2) accounted for 76.8% of the total variance (**Fig. 1D**). To account for the uneven number of syllables across behavioral categories, we performed a bootstrap test after downsampling to match the smallest category’s size. This analysis confirmed significant differences in acoustic characteristics between the IntCed and ResCing groups, as well as between the ICing and ResCing groups (IntCed vs ResCing: *p* < 0.001, IntCing vs ResCing: *p*<0.001, IntCed vs IntCing: *p* = 0.510; **Fig. 1E**). Examination of the factor loadings indicated that meanfreq and maxfreq were the primary features contributing to the observed acoustic divergence between these behavioral contexts. Thus, these analyses revealed that the behavioral context of male mice, particularly during IntCed and ResCing interactions, is associated with the production of acoustically distinct USVs.

### 3.2. Specific USV syllables elicit approach behavior in a novel environment

In behavioral experiments, habituation to an apparatus is commonly employed to ensure stimulus-specific attention. While effective, this procedure can raise the motivational threshold for initiating exploratory behavior, potentially leading to an underestimation of behaviors driven by weaker motivation. We hypothesized that conducting playback experiments in a novel environment would maximize the detectability of approach behavior by combining exploration directed toward the apparatus with innate responses to USVs. To test this, we compared the responses of naive mice to USV playback and control stimuli.

Auditory stimuli were sourced from USVs recorded during IntCed and ResCing. Representative syllable segments were selected for playback (**Fig. 2A, Supplemental Fig. 2**), with PCA (**Fig. 2B**) and Mann–Whitney U test (**Supplemental Fig. 3**) confirming significant acoustic separation between the two groups along the PC1 and PC2 axes. Playback experiments were conducted in an I-maze (**Fig. 2C**), specifically designed to accommodate the high directivity of the ultrasonic sound emitted by the speaker (**Supplemental Fig. 4**). This linear arrangement was adopted to ensure a more uniform sound pressure level along the animal’s approach path and to reduce the spatial bias that may occur in the open field or three chambers commonly used in other studies. Once male mice (n = 18) voluntarily remained at one end of the I-maze, auditory stimuli were delivered from the opposite end. This setup allowed quantification of both approach behavior toward the sound source (**Fig. 2D**) and overall exploratory activity (**Fig. 2E**). Regarding exploratory behavior, no significant differences were observed between USVs and noised conditions for either stimulus type (Wilcoxon signed-rank test; IntCed; W = 79.0, *p* = 0.777 and ResCing; W = 75.5, *p* = 0.711). Whereas the latency to approach the sound source was evaluated using Kaplan–Meier survival curves and the log-rank test (**Fig. 2F**). In the IntCed group, approach latency was significantly shorter during USVs playback than during the noised control (*p* = 0.018). In contrast, no significant difference was observed in the ResCing group (*p* = 0.140), indicating that the attractive effect of USVs is context-dependent. To further examine the temporal consistency of these responses, we quantified synchrony in the distance to the sound source during the first 15 seconds of playback, when all mice had entered the speaker zone, by calculating pairwise correlations across individuals (**Fig. 3A, C**). The distribution of correlation coefficients—calculated among subjects within each group—showed a highly significant difference between the USVs and noised conditions for the IntCed group (*p* = 0.0001; **Fig. 3B**), whereas no such difference was observed for the ResCing group (*p* = 0.764; **Fig. 3D**). This result confirms that the approach behavior triggered by IntCed-derived USVs is not only faster but also characterized by a highly stereotyped temporal pattern that is consistent across individuals.

**Figure 3.**
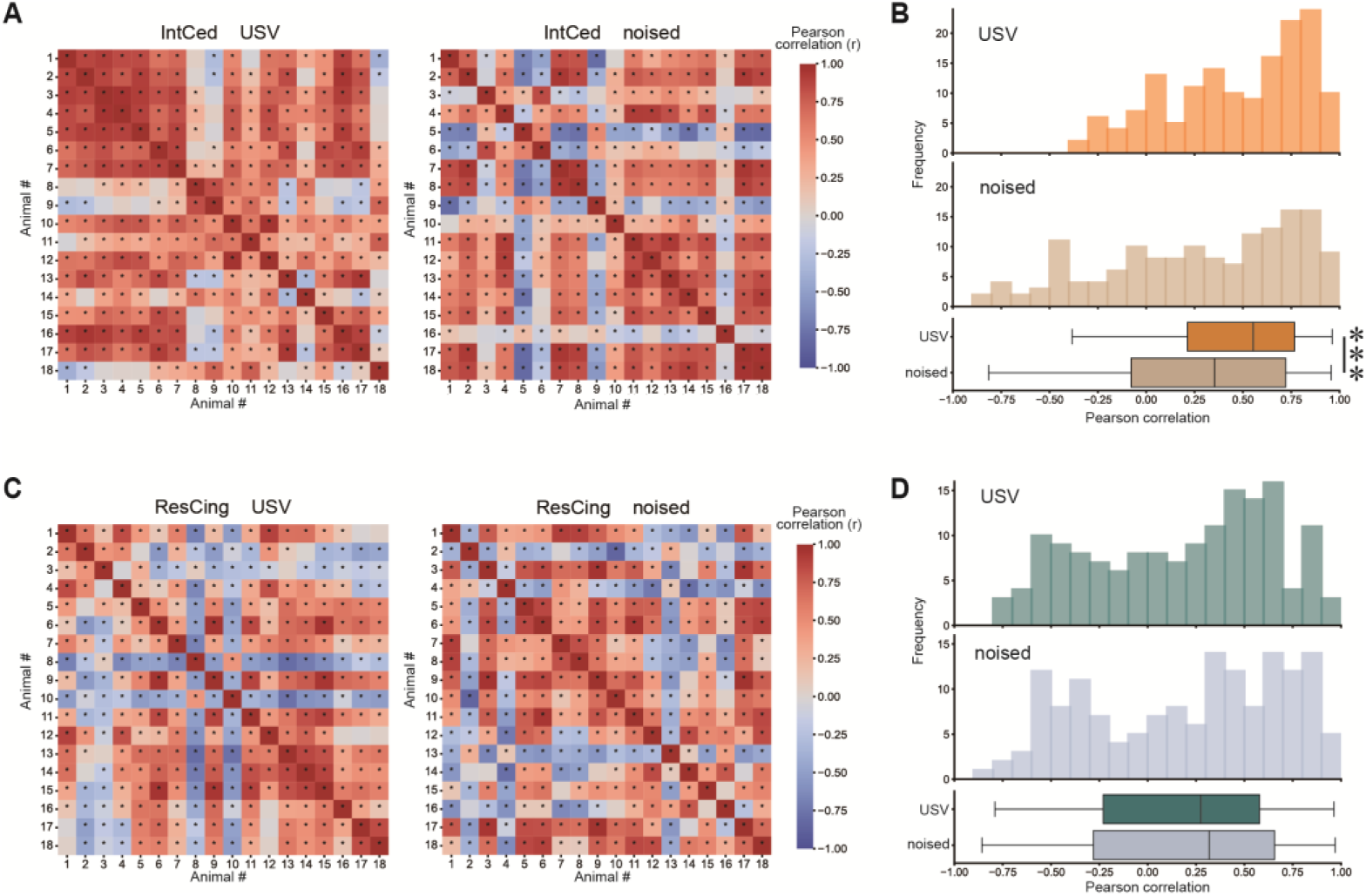
High inter-individual consistency in approach patterns toward chased-intruder (IntCed) USVs. (A) Correlation matrices of approach trajectories (0–15 s) for IntCed. *: *p*<0.05, Pearson correlation coefficient. (B) Comparison of correlation coefficient distributions (top, histogram; bottom, box plot) between USV and noised playbacks in IntCed. ***: *p*<0.001, Wilcoxon signed-rank test. (C) Correlation matrices of approach trajectories (0–15 s) for ResCing. *: *p*<0.05, Pearson correlation coefficient. (D) Comparison of correlation coefficient distributions (top, histogram; bottom, box plot) between USV and noised playbacks in ResCing.

### 3.3. Specific USVs promote exploratory behavior in mice habituated to the apparatus

Another group of male mice (n = 9) acclimated to the apparatus was observed during sound playback (**Fig. 4A**). In contrast to the results observed in a novel environment, exploratory behavior was significantly modulated by the playback of IntCed USVs (χ² = 8.40, *p* = 0.015; **Fig. 4B**). However, no significant change in approach latency was detected (IntCed: *p* = 0.308, ResCing: *p* = 0.186; **Fig. 4C**). Subsequent post-hoc analysis with Holm’s correction revealed a significant increase in exploratory activity during the playback of USVs compared to the noised control (USVs vs noised: *p* =0.023, USVs vs silent: *p*=0.258, noised vs silent: *p*=0.779), while no significant modulation of exploratory behavior was observed for the ResCing group (χ² = 0.22, *p* = 0.895; **Fig. 4C**). To further examine whether synchronized responses occurred despite the absence of significant changes in approach latency, the distance to the sound source during the first 15 seconds after playback onset was analyzed using the same method as in the non-habituated condition. The distribution of all-to-all correlation coefficients among subjects revealed a significant difference between IntCed USVs and noised conditions (*p* < 0.001; **Fig. 5A, B**), whereas no such difference was found for ResCing (*p* = 0.540; **Fig. 5C, D**). The observation of temporal consistency in the IntCed group, even though the approach latency was not fast, suggests that IntCed syllables may be associated with stereotyped behavioral responses. Further analysis revealed no difference between approach and withdrawal velocities in the non-habituated condition, whereas the habituated group exhibited significantly faster approach velocities compared to their withdrawal velocities (**Supplemental Fig. 5**). These results indicate that while approaching the speaker in a novel environment is primarily part of general contextual exploration, the approach behavior in the habituated condition represents a clear, targeted response to the acoustic stimulus, thereby strongly supporting our hypothesis.

**Figure 4.**
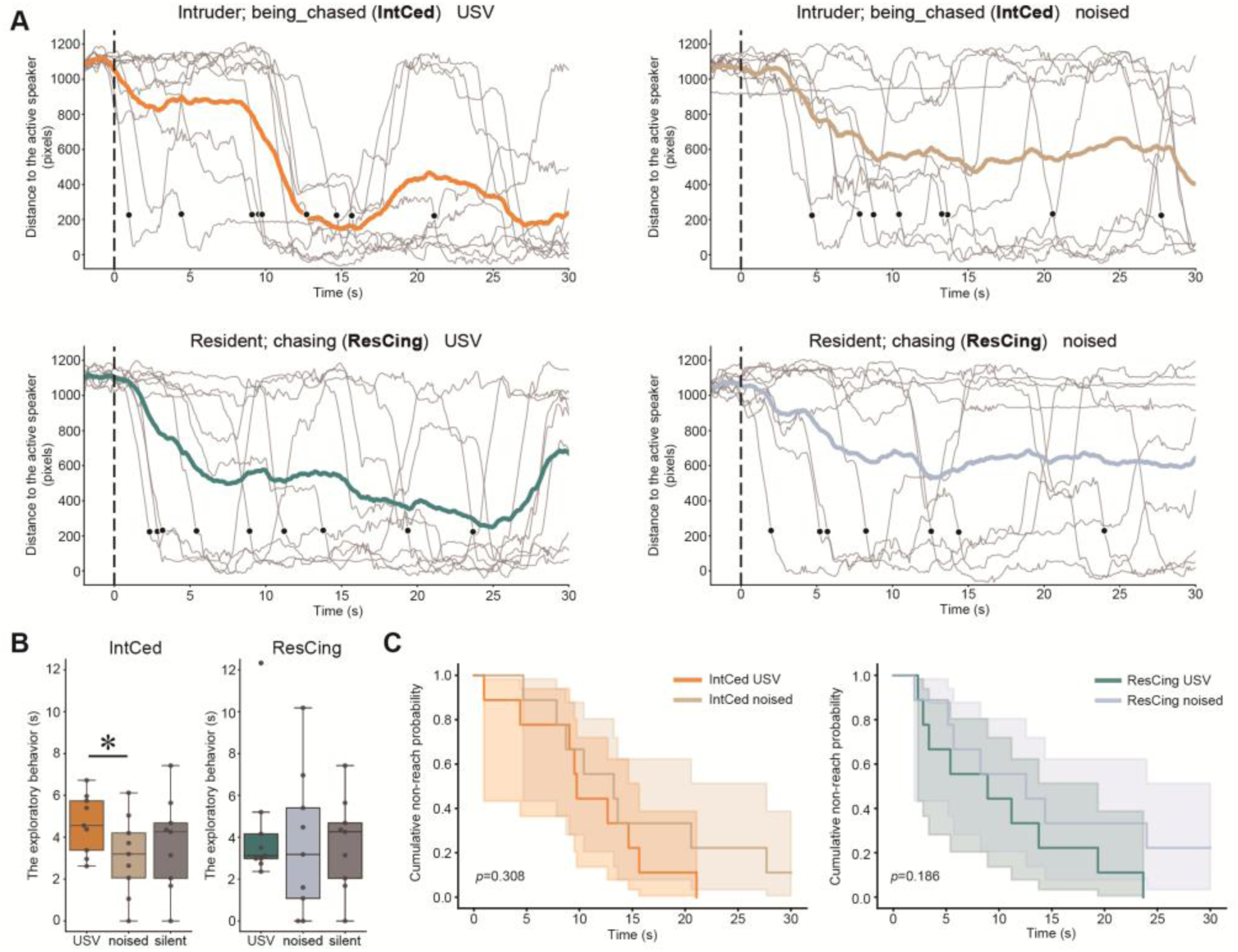
Male mice explore for the source of USVs emitted by chased intruders in a habituation context. (A) Time course of the distance to the active speaker. Gray lines represent individual data from 9 mice, and bold-colored lines represent the mean. The dashed line indicates the onset of the auditory stimulus, and the black dot marks the first entry into the speaker zone. (B) Comparison of exploratory behavior toward the active speaker among USV, noised, and silent conditions in the IntCed (left) and ResCing (right) groups. *: *p*<0.05, Friedman test followed by Holm’s post hoc test. (C) Survival analysis of time to enter the active speaker zone. The shaded area indicates 95% CI.

**Figure 5.**
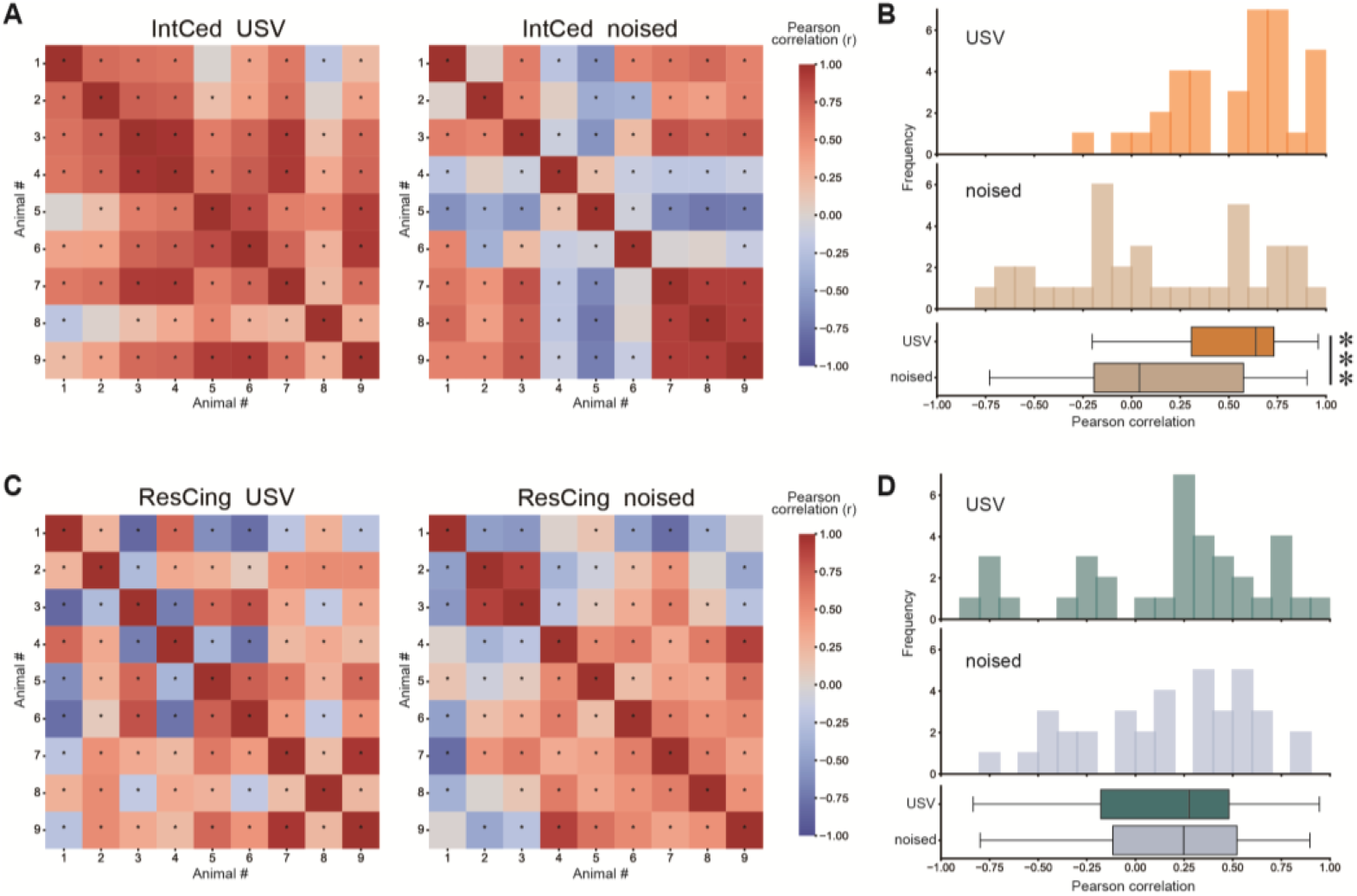
High inter-individual consistency in approach patterns toward chased-intruder (IntCed) emitted USVs. (A) Correlation matrices of approach trajectories (0–15 s) for IntCed. *: *p*<0.05, Pearson correlation coefficient. (B) Comparison of correlation coefficient distributions (top, histogram; bottom, box plot) between USV and noised playbacks in IntCed. ***: *p*<0.001, Wilcoxon signed-rank test. (C) Correlation matrices of approach trajectories (0–15 s) for ResCing. *: *p*<0.05, Pearson correlation coefficient. (D) Comparison of correlation coefficient distributions (top, histogram; bottom, box plot) between USV and noised playbacks in ResCing.

### 3.4. Upward frequency-modulated syllables induce approach behavior in male mice

Finally, to identify the specific vocalizations that trigger approach behavior, we defined a period within the playback stimuli that consistently elicited approach behavior. We calculated all-to-all correlation coefficients for distance to the sound source within 1-second windows. An approach event was defined as a period during which the correlation coefficient was significantly above the chance level, and the mice were actively moving toward the sound source. In the IntCed group, approach events were consistently defined during the 0–5 s interval in the novel context (**Fig. 6A**) and the 7–12 s interval in the familiar context (**Fig. 6B**). We also analyzed syllables from the 0–5 s interval of ResCing USV playback as a behaviorally ineffective condition, because no approach behavior was observed despite the mice having sufficient spatial opportunity to move toward the sound source. Examination of the spectrograms from these three sections revealed differences in syllable composition. In the novel environment, IntCed USV sequences were predominantly composed of up-fm syllables (dots in **Fig. 6C**). Likewise, in the familiar environment, the approach behavior was also associated with a high prevalence of up-fm syllables. In contrast, ResCing USV sequences were characterized by a markedly different acoustic profile, with a lower proportion of up-fm syllables (**Fig. 6C**). Next, to determine whether the acoustic characteristics of syllables differed among these three groups, we performed PCA using five acoustic parameters, as in **Fig. 1D**. We then compared the distributions of syllables by quantifying within-group and between-group distances in the PCA space. This analysis revealed that the between-group distances between both novel and familiar IntCed USVs and ResCing USVs were significantly larger than the corresponding within-group distances (novel IntCed vs ResCing: *p* < 0.001, familiar IntCed vs ResCing: *p* < 0.001, novel IntCed vs familiar IntCed: *p* = 0.118; **Fig. 6D**). Specific parameters driving this separation were identified by focusing on mean frequency and maximum amplitude. A Kruskal–Wallis test revealed significant differences across the groups for both parameters (H = 14.337, *p* < 0.001; **Fig. 6E, H** = 24.268, *p* < 0.001; **Fig. 6F**). Subsequent post-hoc multiple comparisons confirmed that syllables in both the IntCed Novel and IntCed Familiar groups had significantly higher mean frequencies (novel IntCed vs ResCing: *p* < 0.001, familiar IntCed vs ResCing: *p* = 0.007; **Fig. 6E**) and maximum amplitudes (novel IntCed vs ResCing: *p* = 0.007, familiar IntCed vs ResCing: *p* < 0.001; **Fig. 6F**) compared to those in the ResCing group, mirroring the structural differences observed in the PCA. These findings suggest that the specific acoustic profile of IntCed USVs—characterized by higher pitch and greater sound pressure—has greater acoustic salience, which may effectively trigger approach behavior.

**Figure 6.**
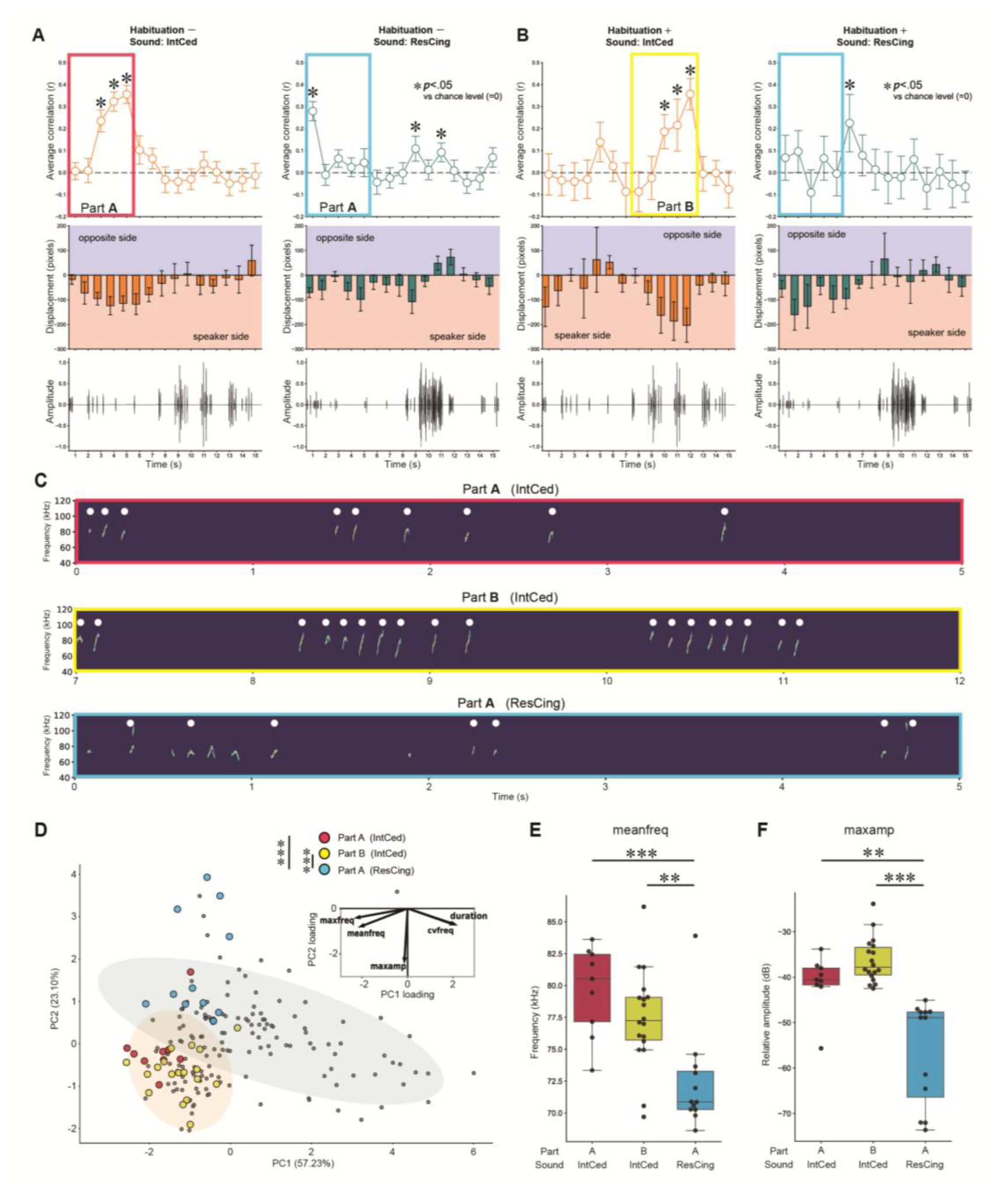
USV syllables eliciting male mice approach exhibit specific acoustic characteristics. (A) Relationship between mouse movement and timing of syllables in a non-habituation context. The red-outlined area indicates the time window defined as approach behavior, and the light blue square represents its control. (Top) Correlation coefficient for distance to speaker. (one-sample t-test; \**p* < 0.05 vs. chance level of 0) (Middle) Movement distance to the speaker. (Bottom) Amplitude of the presented sound stimuli. (B) Relationship between mouse movement and timing of syllables in a habituation context. The yellow-outlined area indicates the time window defined as approach behavior. (Top) Correlation coefficient for distance to speaker. *: *p*<0.05, one-sample t-test vs. chance level of 0. (Middle) Movement distance to the speaker. (Bottom) Amplitude of the presented sound stimulus. (C) Spectrogram of USVs during the time window indicated by the squares in (A) and (B). The syllables marked with a white dot indicate those classified as Up-FM. (D) Scatter plot of syllables emitted in the time windows of panels (A) and (B) (color circles) and in the rest of the playback (gray dots) plotted on PC axes of the acoustic features of USVs used in the playback experiment. ***: *p*<0.001, Mann-Whitney U test followed by Holm’s post hoc test. (E) Mean frequency of syllables contained in each time window. ***: *p*<0.001, **: p<0.01, Kruskal–Wallis test followed by Dunn test. (F) Max amplitude of syllables contained in each time window. ***: *p*<0.001, **: p<0.01, Kruskal–Wallis test followed by Dunn test.

## 4. Discussion

Our findings demonstrate that male mice emit distinct types of USVs depending on their social role—whether they are a resident that is chasing an intruder or an intruder that is being chased by a resident—and that these vocalizations possess different communicative values. Specifically, playback experiments revealed that USVs emitted during being chased (IntCed USVs) elicit approach behavior in receiver mice. Notably, this response showed high temporal synchrony across subjects, a phenomenon that persisted regardless of the mice’s prior acclimation to the apparatus. The acoustic analysis associated with this behavior identified upward-frequency-modulated syllables, with higher mean frequency and greater sound pressure, as a key factor in this exploratory approach. Together, these results suggest that certain USV structural features serve as potent social signals that trigger stereotyped behavioral responses in a context-independent manner. Importantly, similar approach tendencies toward IntCed USVs were observed in an open-field arena during preliminary trials (**Supplemental Fig. 6, 7**), suggesting that the response is not an artifact of the specific maze geometry, even though the I-maze was employed to ensure the acoustic precision necessary for rigorous evaluation.

### 4.1. Distinct acoustic features of USV in different social contexts

USVs emitted during IntCed and ResCing were composed of syllables with overall distinct acoustic characteristics. Traditionally, syllable types have been classified based on spectrogram shapes using manual labeling or machine-learning approaches, often relying on study-specific criteria that can be inherently arbitrary. While dimensionality reduction techniques have been implemented to mitigate this limitation, clearly distinguishing individual syllables in mice remains challenging compared to songbirds, as they often do not form discrete clusters^27^. Therefore, we investigated whether the acoustic characteristics of syllables differ across entire USV sequences as a function of social context. Our results demonstrated that IntCed and ResCing USVs exhibit distinct syllable compositions, a pattern consistently observed across both classical spectrogram-based classification and dimensionality-reduction–based analyses. In particular, the higher prevalence of up-fm syllables in the vocalizations of being chased individuals is consistent with previous reports ^20,21^. Moreover, a study in female mice ^21^ reported that individuals emitting USVs in aggressive contexts tended to engage in chasing behavior, whereas those emitting USVs in non-aggressive contexts tended to exhibit withdrawal behavior; notably, the latter exhibited a higher mean frequency, consistent with this study (**Fig. 1E and 6E**). These results reinforce the validity of analyzing USVs by segmenting them according to specific social behaviors. The present work suggests consistency in mouse vocal communication, highlighting the functional importance of these acoustic features in mediating social interactions.

### 4.2. Functional significance of male-attracting USVs

Male mice exhibited approach behavior toward IntCed USVs but not toward ResCing USVs; however, the underlying reason remains unclear. Considering the known relationships between acoustic structure and social intent, this behavior may be explained through two complementary perspectives. First, while up-fm syllables are stably prevalent across mouse strains ^28–30^, their proportion decreases during active social interactions such as sniffing or chasing, as observed in our results (**Supplemental Table 1**) and a previous study ^31^. This suggests that the syllables which increase in these contexts (e.g., chevrons or frequency steps) may function as signals of the caller’s own active approach rather than as cues to attract a receiver. Second, the functional significance of IntCed vocalizations can be inferred from studies in other rodents. In rats, 50-kHz calls are thought to prevent male-male play from escalating into aggression, and devocalization has been shown to increase aggressive encounters ^32^. Similarly, gerbils emit high-frequency vocalizations, including bent up-fm, when they show non-agonistic responses to aggressive encounters, and a higher occurrence of these calls is associated with a reduced likelihood of further aggression ^33^. Such vocalizations are often considered contact calls, and playback experiments have demonstrated that conspecifics approach their sound source ^9,34^. In mice, previous research also shows that non-jumping high-frequency syllables decrease as aggression induced by early-life stress increases ^35^, suggesting a strong association between exposure to aggression and specific USVs in rodents. Taken together, the up-fm calls emitted by the intruder may represent a kind of contact call that communicates non-hostility and facilitates the establishment of social relationships, thereby promoting the observed approach behavior in listener mice.

### 4.3. Temporal organization of USV syllables

Previous research has demonstrated that mouse USVs possess a characteristic rhythmic structure within these bouts ^36^. Notably, when male songs were presented to females, disrupting this rhythm led to a marked reduction in approach behavior relative to the original, indicating that syllable-to-syllable timing is critical ^14^. In addition, randomizing the syllable order within a sequence significantly decreases the approach response compared to the natural sequence ^37^. Therefore, when mice are habituated to the environment and their exploratory motivation is reduced, the temporal structure of these bouts may provide additional cues that help them reach the behavioral threshold for approach. Collectively, these findings indicate that USV-driven behavior is shaped not only by the properties of individual syllables but also by their temporal organization.

Overall, our findings show that male mice produce distinct USVs depending on their social role and that only vocalizations of chased intruders reliably induce approach behavior. This response appears to depend on both the acoustic features of individual syllables and their temporal organization. These results highlight how social context shapes vocal output and how listeners use multiple levels of acoustic information to guide social decisions.

### 4.4. Limitations of the study

Our study has several limitations to consider. First, our behavioral paradigm relied exclusively on auditory playback, whereas mouse social communication is inherently multimodal. Given that olfactory cues play an important role in social interaction and can modulate responses to USVs ^11,38^, future investigations should examine how integrating vocal and olfactory information influences the salience of these signals. Second, the experimental setup was designed to assess approach behavior, thereby precluding the observation of potential avoidance or aversive responses. To determine whether certain USVs actively induce avoidance rather than simply failing to attract, paradigms that deliver sound stimuli in close proximity to the animal should be employed. Indeed, our preliminary experiment using a different type of behavioral arena produced avoidance behavior in ResCing USVs. Finally, the neurophysiological mechanisms underlying the observed approach behavior remain unknown. Specifically, given that pup calls trigger maternal approach behavior while reflecting a negative emotional state in the receiver ^12^, the elicitation of approach behavior does not inherently signify that the underlying USVs represent a “positive” signal. Future studies incorporating in vivo physiological recordings in freely moving mice will be necessary to identify the neural circuits that detect these USVs and to clarify the emotional processes that drive the distinct behavioral outcomes.

---

**Supplementary Table 1.** Residual Analysis of Syllable Proportions in 3 Different Social Conditions. Residuals with absolute values larger than 3 are shown in bold, indicating strong deviation.

|  | up fm | down fm | flat | short | chevron | step-up | step-down | two-steps |
| --- | --- | --- | --- | --- | --- | --- | --- | --- |
| ICed | **5.628** | -1.85 | -0.505 | -1.386 | **-3.265** | -1.218 | 0.626 | -2.144 |
| ICing | 2.164 | -0.121 | -0.447 | -0.414 | -0.803 | -1.003 | -1.866 | 0.31 |
| RCing | **-6.32** | 1.642 | 0.753 | 1.468 | **3.332** | 1.762 | 0.851 | 1.571 |

**Supplementary Figure 1.**
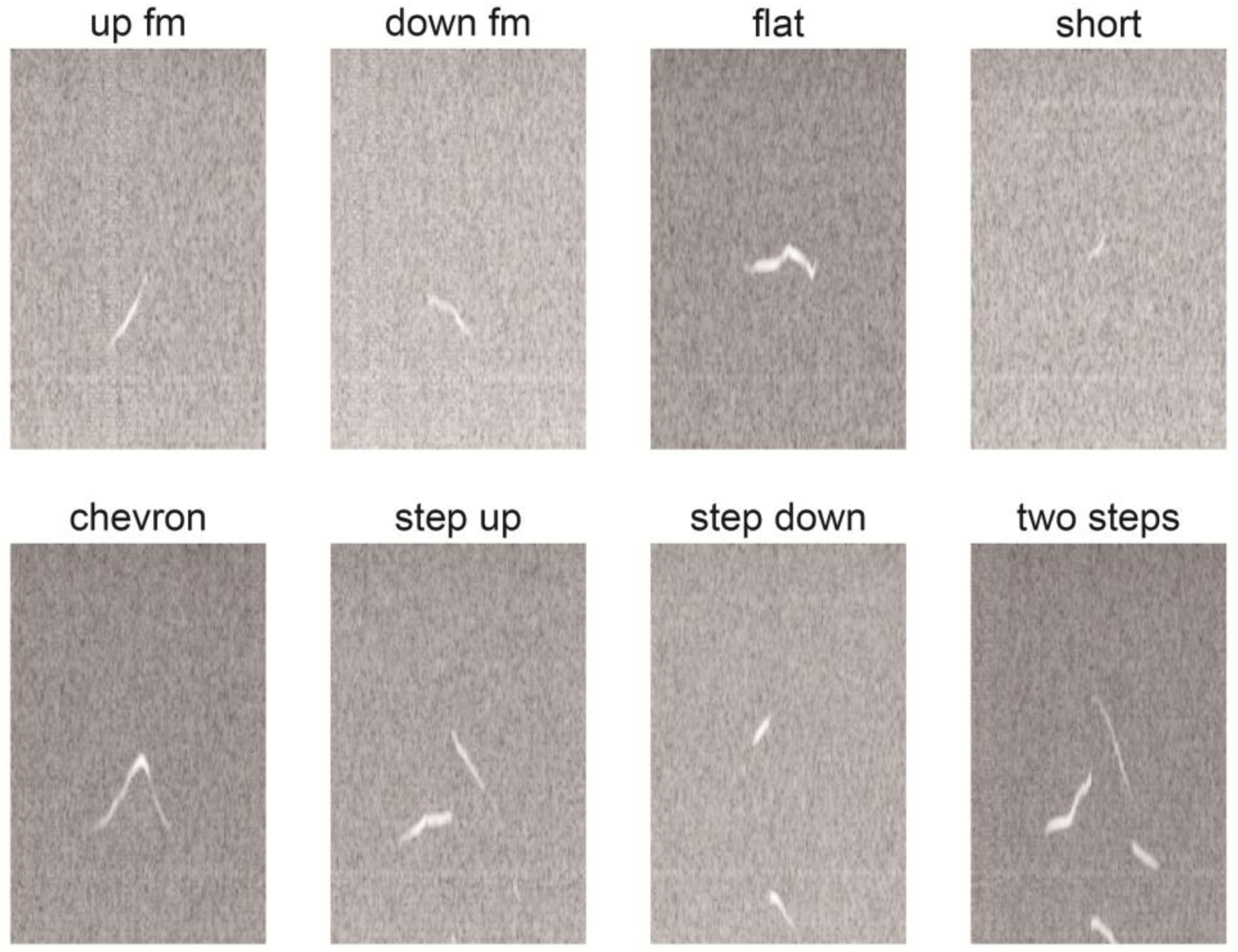
Representative spectrograms of the eight call types identified in USVs recorded during male-male interactions.

**Supplementary Figure 2.**
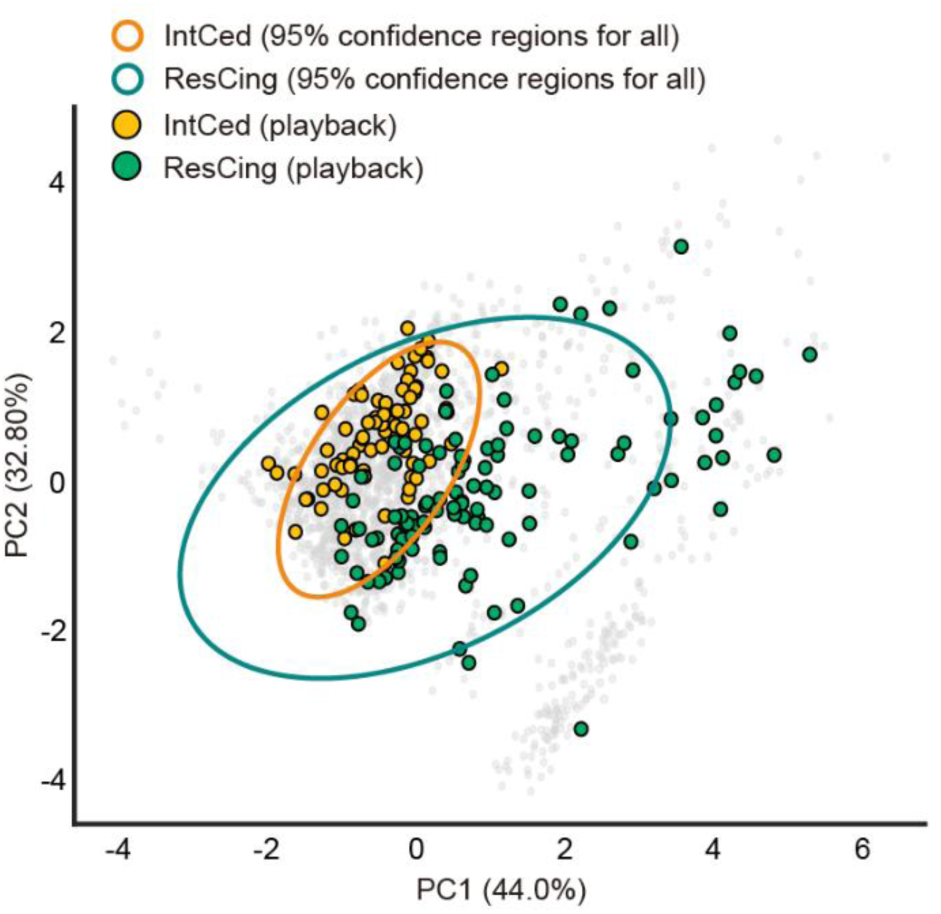
Distribution of syllables selected for playback experiments within the 2D PCA space of all syllables. Gray dots represent the entire population of detected syllables, with ellipses indicating the 95% confidence regions for IntCed (orange) and ResCing (green) groups. Filled circles highlight the specific syllable segments selected for playback experiments (IntCed, yellow; ResCing, lightgreen).

**Supplementary Figure 3.**
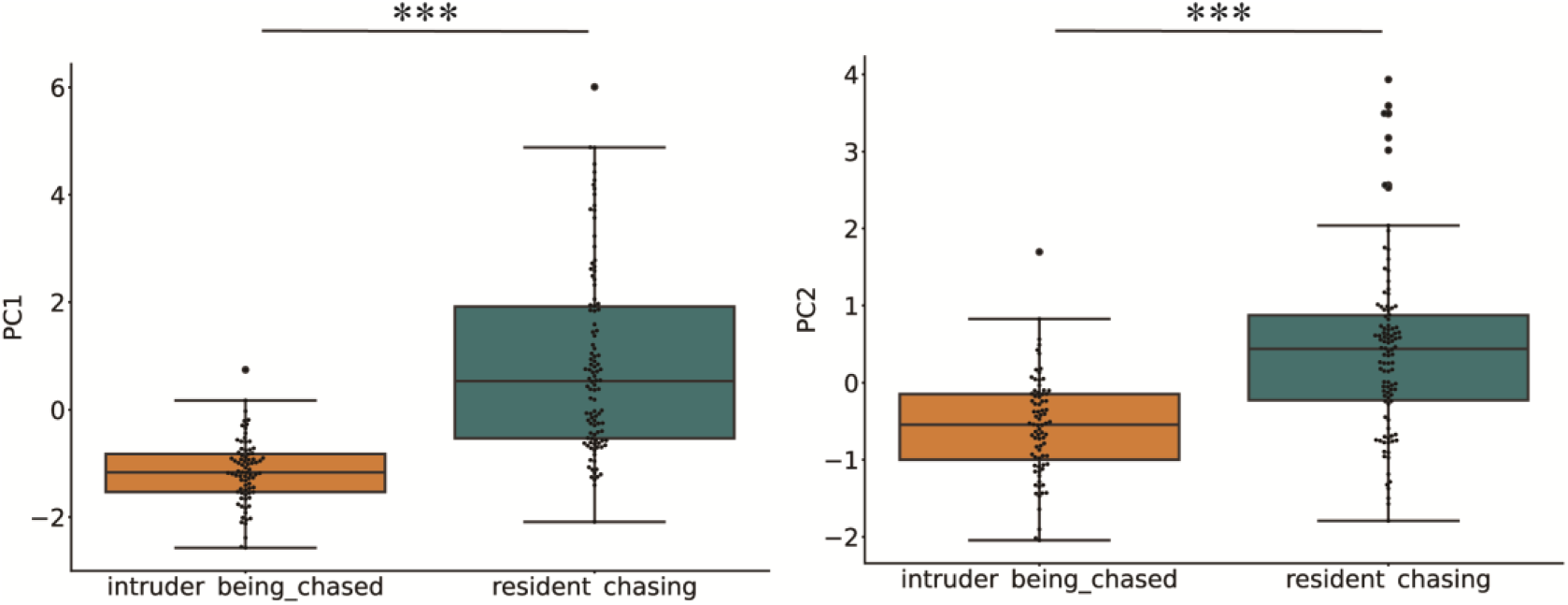
The Acoustic Characteristics of IntCed and ResCing USVs Differ. Bar graphs of two PCs of the acoustic characteristics of the USVs used in the playback experiment show a significant difference between the social contexts. ***: *p*<0.001, Mann-Whitney U test.

**Supplementary Figure 4.**
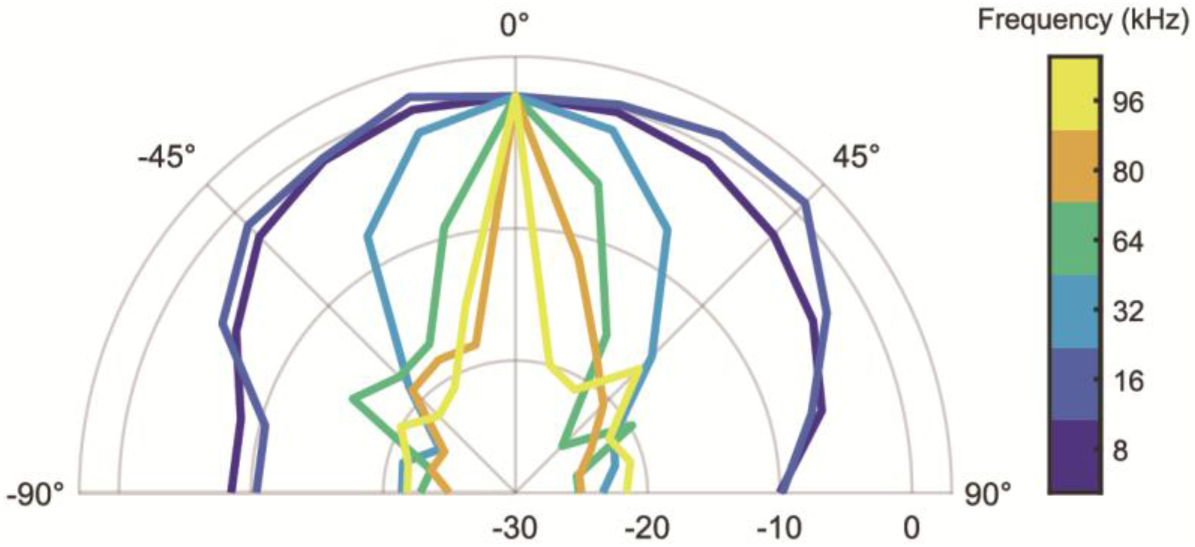
The speaker (ES1, TDT) directivity by frequency.

**Supplementary Figure 5.**
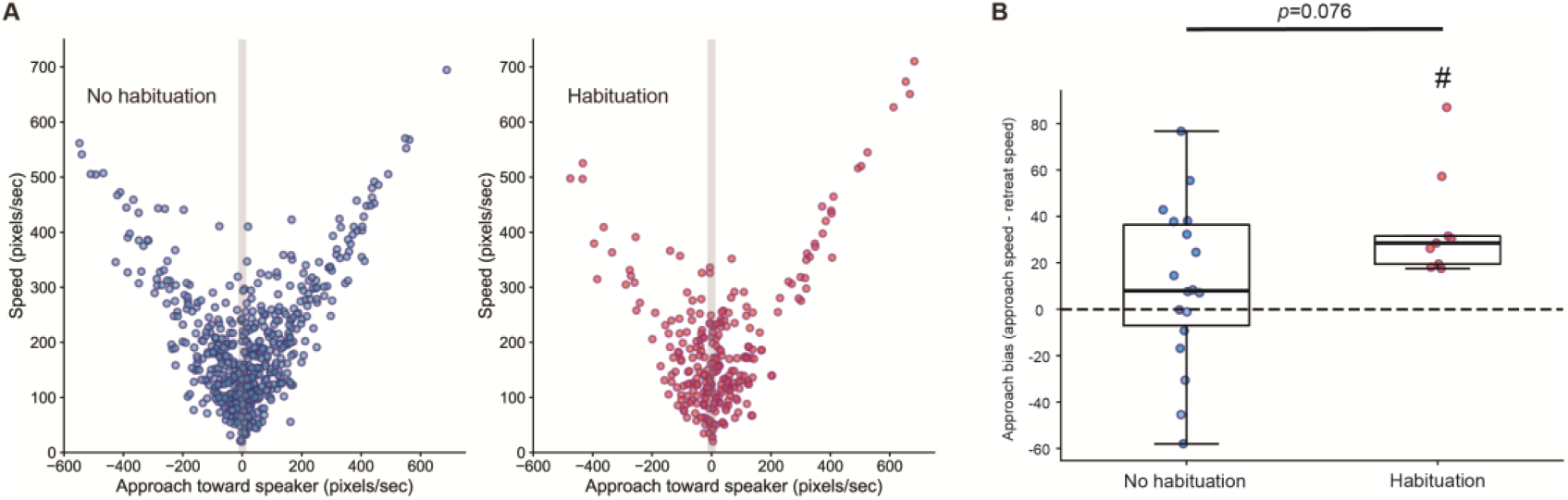
Habituation increases approach-biased locomotion toward the speaker. (A) Scatter plots showing the relationship between velocity and the change in distance to the speaker for each second of the playback experiments (IntCed USV). Positive values on the x-axis indicate movement toward the speaker (approach), whereas negative values indicate movement away from the speaker (retreat). Data are shown for the no-habituation (left) and habituation (right) conditions. The vertical gray band indicates a threshold region (±10 pixels/sec) introduced to exclude small positional fluctuations due to tracking noise. Data points within this range were excluded from subsequent analyses. (B) Approach bias calculated for each animal as the difference between the mean velocity during the approach and retreat periods. Each dot represents one animal, and box plots show the median and interquartile range. The dashed line indicates zero bias. In the habituation group, the approach velocity was significantly higher than the withdrawal velocity (#: *p*<0.01 vs. chance level of 0, Wilcoxon signed-rank test.) and also tended to be higher than the approach velocity observed in the no-habituation group. Mann–Whitney U test, U = 46, *p* = 0.076; effect size *r* = 0.43, rank-biserial correlation.

**Supplementary Figure 6.**
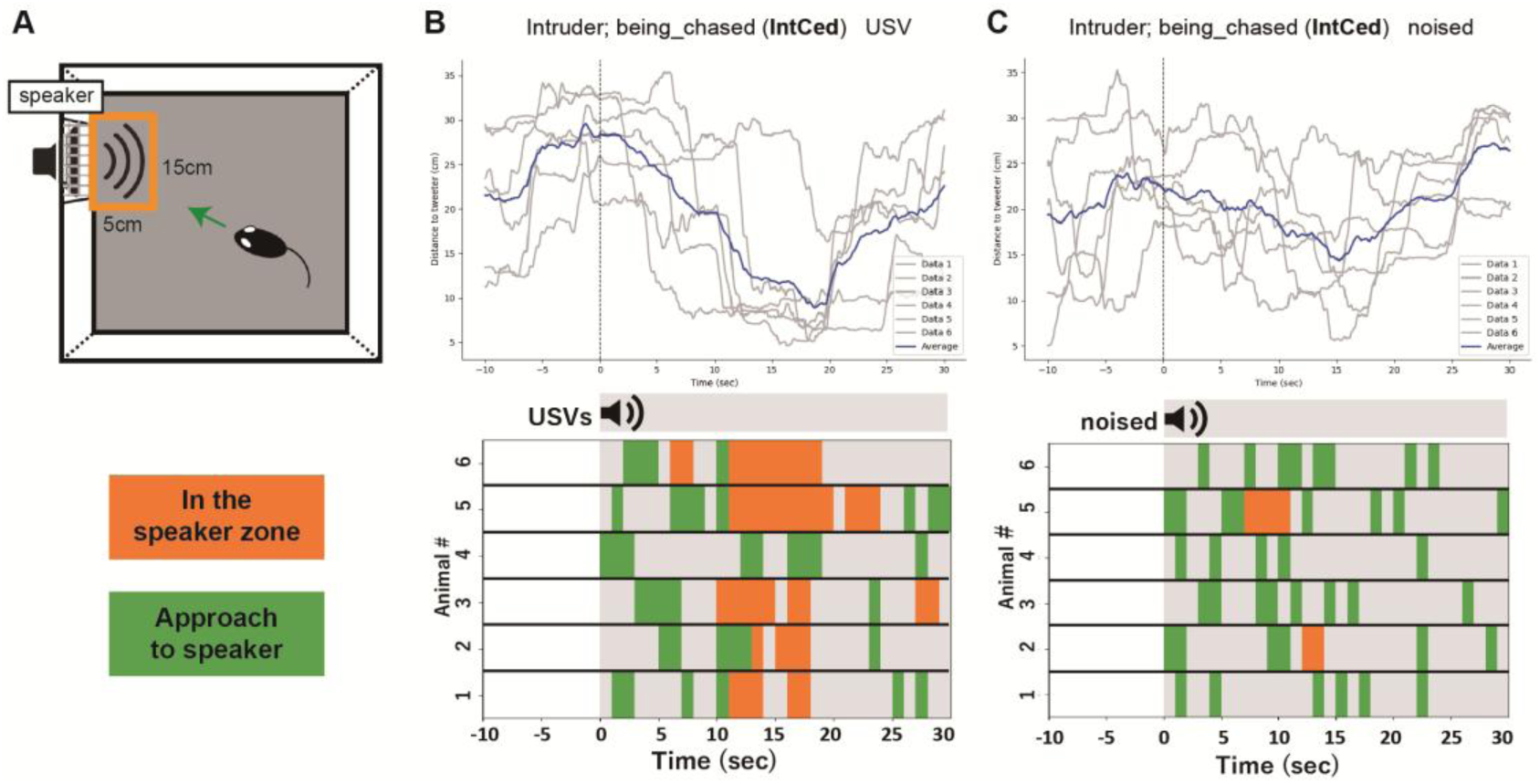
Results of Playback Experiments in Open Fields (IntCed) The behavior of male mice in response to USVs and noised was observed (Brightness 10 lux), using an open field (30cm×30cm) with mesh holes in the sides. After a 5-minute habituation period to the apparatus, mice were presented with 60 seconds of ultrasonic or noised (70-75 dB SPL) when they were not in the area in front of the speaker (15 cm × 15 cm). The following day, the same experiment was conducted under different conditions. (A) Schematic diagram of the open field test for Playback Experiments. The orange square is defined as the speaker zone. (B) The behavior when presented with IntCed “USV”. (Top) Time course of distance to speaker. (bottom) Color-coded behavioral outcome in response to sound stimulus in 6 mice. (C) The behavior when presented with IntCed “noised”. (Top) Time course of distance to speaker. (bottom) Color-coded behavioral outcome in response to sound stimulus in 6 mice.

**Supplementary Figure 7.**
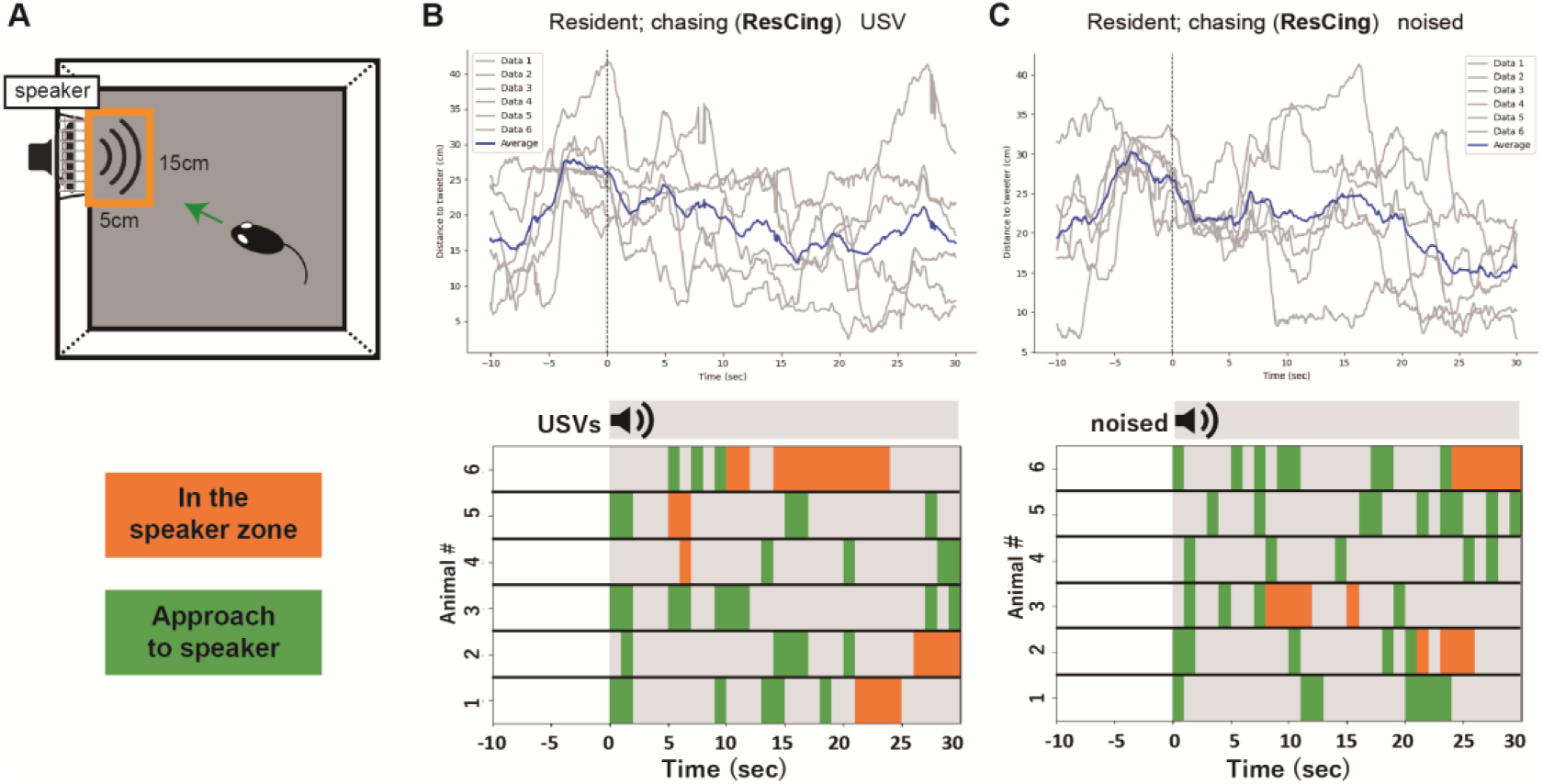
Results of Playback Experiments in Open Fields (ResCing) (A) Schematic diagram of the open field test for Playback Experiments. The orange square is defined as the speaker zone. (B) The behavior when presented with ResCing “USV”. (Top) Time course of distance to speaker. (bottom) Color-coded behavioral outcome in response to sound stimulus in 6 mice. (C) The behavior when presented with ResCing “noised”. (Top) Time course of distance to speaker. (bottom) Color-coded behavioral outcome in response to sound stimulus in 6 mice.

